# Peripheral Axons Form Endoplasmic Reticulum-Rich Conduits With Breast Cancer Cells

**DOI:** 10.64898/2026.11.02.763595

**Authors:** Neşe Ayşit, Şükriye Bilir, Esra Altıntaş, Fatih Toy, Mehmet Şerif Aydın, Hasan Korkaya, Gürkan Öztürk

## Abstract

Peripheral nerves are active components of the tumor microenvironment, but the physical routes by which sensory axons communicate with cancer cells remain incompletely defined. Here, we show that primary dorsal root ganglion neurons form thin membrane conduits with EMT6 breast cancer cells that contain endoplasmic reticulum (ER). ER-Tracker labeling and neuron-restricted expression of the ER proteins CNPY2 and ATL3 identified ER-rich axonal extensions terminating on cancer cells. Serial-section scanning transmission electron microscopy resolved ER-like membranes within these conduits and at axon-cancer cell interfaces. Scanning electron microscopy and live holotomographic imaging further revealed dynamic contact formation and associated remodeling of the cancer cell surface. These observations define an ER-containing structural interface between sensory axons and cancer cells and provide a framework for testing whether neuronal ER contributes to direct intercellular signaling or material transfer.

**Summary:** Aysit et al show that primary sensory axons form thin endoplasmic reticulum-containing conduits with breast cancer cells. Fluorescence, electron microscopy and live holotomography reveal ER-rich neuronal contact domains, defining a structural route for a direct neuron-tumor interaction.

## Introduction

Peripheral nerves are increasingly recognized as active components of solid-tumor microenvironments, where they can influence tumor growth, invasion, and metastasis (Aysit et al., 2026; Fu et al., 2026; Jiang et al., 2026; Winkler et al., 2026). Beyond diffusible neurotransmitter signaling, neurons can communicate directly with cancer cells through synapse-like contacts or membrane conduits, and nerve-to-cancer mitochondrial transfer has recently emerged as one form of such communication (Hoover et al., 2025; Ren et al., 2025). Tunneling nanotubes (TNTs) and related membrane protrusions can support intercellular exchange of organelles and signaling components, but their structures and modes of connectivity are heterogeneous (Cordero Cervantes and Zurzolo, 2021; Yamada et al., 2026). Axons contain a continuous tubular endoplasmic reticulum (ER) network with roles in calcium homeostasis, lipid metabolism, and local protein synthesis (Berridge, 1998; Berridge, 2002; Koppers et al., 2024; Ozturk et al., 2020). ER has been reported within intercellular conduits in other cellular contexts (Kadiu and Gendelman, 2011; Wang et al., 2011), but whether axonal ER enters direct neuron-cancer cell connections is unknown. We therefore combined fluorescence labeling, neuron-restricted ER reporters, serial-section electron microscopy, scanning electron microscopy, and live holotomographic imaging in primary dorsal root ganglion (DRG) neuron-EMT6 breast cancer cell co-cultures to determine whether ER is present within axon-derived conduits and their cancer cell contact domains.

## Results

### Sensory axons form ER-labelled contacts with breast cancer cells

Primary DRG neurons from βIII-tubulin-GFP mice were co-cultured with mCherry-expressing murine EMT6 breast cancer cells. Neurons extended axons throughout the cultures while EMT6 cells remained adherent and proliferative. To visualize neuronal ER before cancer cell addition, DRG cultures were labelled with ER-Tracker and extensively washed. Twenty-four hours after EMT6 cells were added, ER-Tracker-positive thin extensions were observed emerging from axons and terminating on cancer cells; we referred to these conduits as TNT-like extensions (TNT-LEs). The terminations were frequently with broadened or foot-like endings (Fig. 1A). In other contacts, ER-Tracker formed a larger cap-like domain at the cancer cell surface (Fig. 1B), whereas en passant axons showed ER-enrichment at points of cancer cell contact (Fig. 1C). These morphologies indicate that axonal contacts with cancer cell can be enriched in ER.

**Figure 1.**
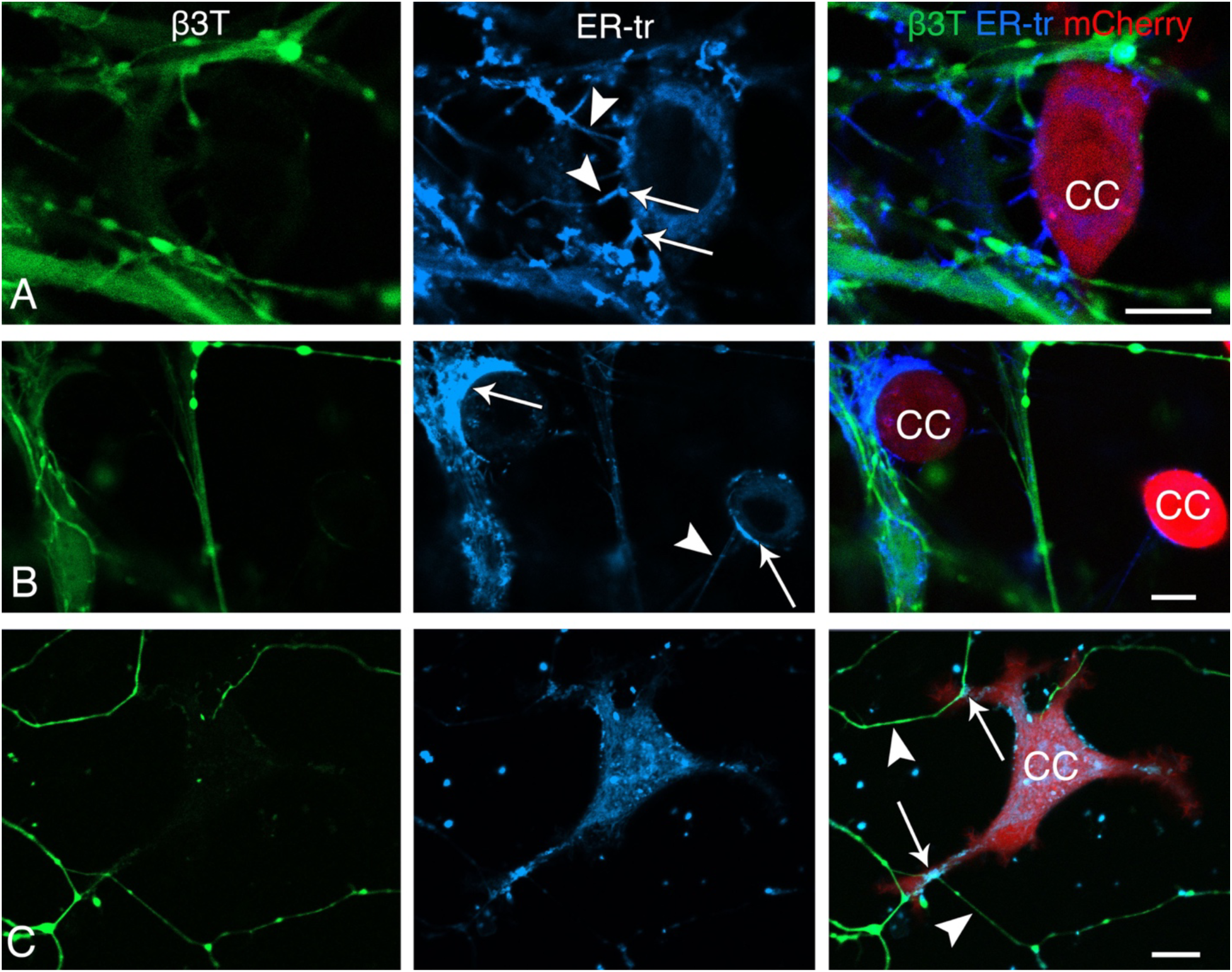
Sensory axons form ER-labeled contacts with EMT6 breast cancer cells. DRG neurons (βIII-tubulin-GFP; green) were labeled with ER-Tracker (blue) before addition of mCherry-EMT6 cells (red). **(A)** ER-Tracker-positive TNT-like extensions (TNT-LEs; arrowheads) emerging from axons and contacting a cancer cell with foot-like terminal enlargements (arrows). **(B)** ER-Tracker-enriched cap-like domain (upper arrow) at a cancer cell surface and a separate ER-labeled TNT-LE (arrow head) with a terminal enlargement (lower arrow). **(C)** En-passant axons (arrow heads) with ER-Tracker enrichment at cancer cell contact sites (arrows). Note that for the purpose of clarity only some of the similar structures are pointed. Scale bars 10 µm. Number of biological replicates (animals) / culture dish per animal / imaged area per culture dish: 3/4/4.

### Axonal ER proteins localize to ER-rich neuron-cancer cell interfaces

Because a small-molecule ER dye cannot by itself establish the cellular origin of labelled membranes, we next expressed GFP-tagged ER proteins selectively in DRG neurons before adding mCherry-EMT6 cells. For this, we selected Atlastin3 (ATL3) and Canopy2 (CNPY2 due to their specific roles in axonal dynamics like neurite growth and ER plasticity (Behrendt et al., 2019; Hatta et al., 2014; Schildknegt et al., 2019). Both constructs labeled axons and thin axon-derived extensions that contacted cancer cells (Fig. 2). Three-dimensional reconstructions showed CNPY2-positive TNT-LEs terminating in discrete ER-rich domains at the cancer cell surface (Fig. 2A and Video 1). ATL3-eGFP produced a similar pattern: ATL3-positive TNT-LEs connected to cancer cells and accumulated at terminal contact domains (Fig. 2 B and Video 2). We refer to these contact-associated structures descriptively as **“E**R-**R**ich **N**euronal **I**nterface **D**omains” (ERNIDs). These experiments support a neuronal origin for the ER signal within the connecting extensions and terminal domains.

**Figure 2.**
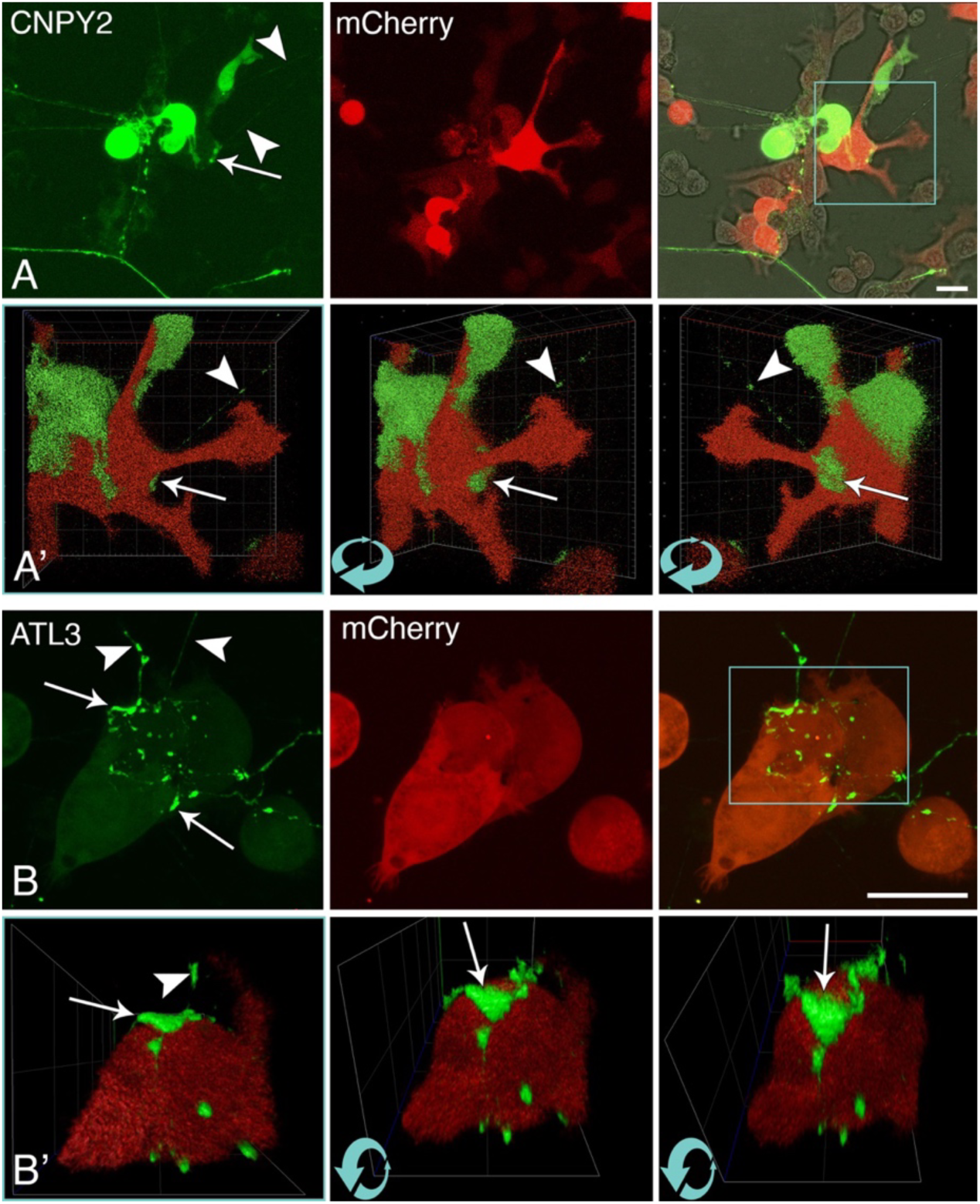
Axonal ER proteins localize to ER-rich neuron-cancer cell interfaces. **(A)** Canopy2 (CNPY2) expression in neurons and in their axons and TNT-LEs (arrow heads) connected to cancer cells (mCherry) with special formations, which we refer to as “ER-rich neuronal interface domain” (ERNID) (arrow). (**A’)** Three-dimensional rotational views of the boxed area in “A” reconstructed from serial optical sections showing a TNT-LE (arrow heads) terminating on cancer cell with an ERNID (arrows). **(B)** Atlastin 3 (ATL3) expression in axons and TNT-LEs (arrow heads) connected to cancer cells with ERNIDs (arrows). **(B’)** Three-dimensional rotational views of the boxed area in “A” reconstructed from serial optical sections showing a TNT-LE (arrow head) terminating on cancer cell with an ERNID (arrows). Note that for the purpose of clarity only some of the similar structures are pointed. Scale bars 20 µm. See also Video 1 and 2. Number of biological replicates (animals) / culture dish per animal / imaged area per culture dish for each protein: 2/4/4.

### Serial-section electron microscopy resolves ER within axon-cancer cell conduits

We next examined the ultrastructure of the axon-cancer cell connections by taking serial sections for scanning transmission electron microscopy (STEM). Thin membrane-bound extensions, TNT-LEs were observed between axons and EMT6 cells (Fig. 3). Within these extensions, elongated membrane profiles with ER-like morphology could be followed across serial sections. In several examples, the ER-containing TNT-LEs seemed to be fused to the cancer cell plasma membrane creating cytoplasmic continuation; in one series, a thin ER profile within the extension lay adjacent to the thicker ER cisterna within cancer cell (Fig. 3). Other interfaces contained clusters of round membrane profiles near contact site. Some extensions terminated within vacuole-like invaginations of cancer cell surface (Fig.S1). Mitochondria were also present in a subset of TNT-LEs, consistent with the broader capacity of neuronal processes to support organelle trafficking (Hooover et a., 2025). However, the present ultrastructural data do not establish transfer of ER or mitochondria across the cancer cell membrane.

**Figure 3.**
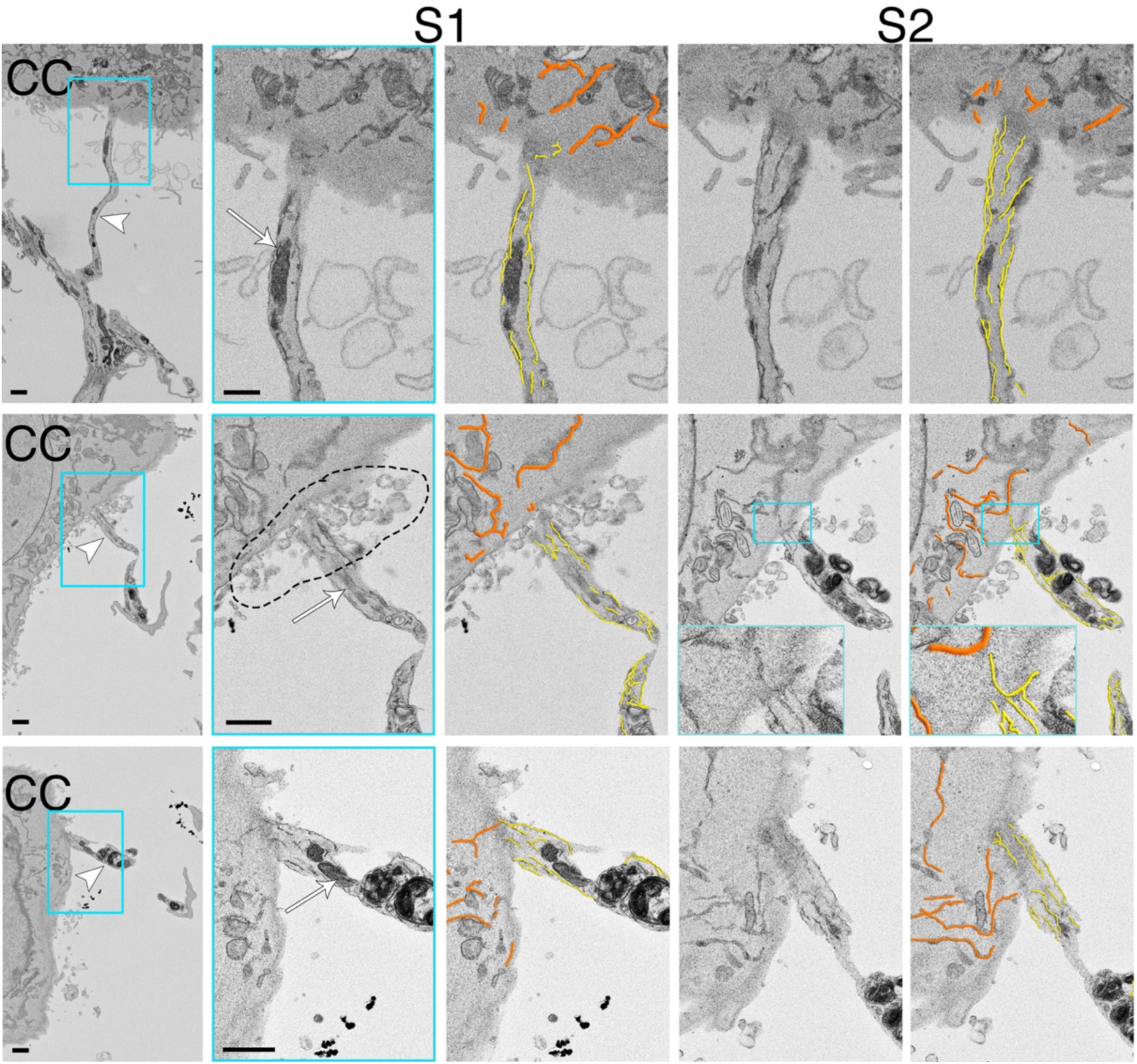
Serial-section STEM resolves ER within axon-cancer cell conduits. Three examples of TNT-LE contact sites between neuronal processes and cancer cells (CC). Boxed regions are enlarged and representative serial sections (S1 and S2) are shown. ER-like membranes are traced in yellow within TNT-LEs and in orange within cancer cells. In the middle example, the inset highlights close apposition of a thin ER profile within the extension and a thicker cancer cell ER profile. The dotted region indicates clustered round membrane profiles near the interface. Scale bars 1 µm. Number of biological replicates (animals) / culture dish per animal / sectioned and imaged area per culture dish: 3/3/1.

### Scanning electron microscopy reveals unique morphological features of axon - cancer cell interfaces

Next, we examined the co-cultures under scanning electron microscope (SEM). There were several TNT-LEs of various diameters between cancer cells and axons or neuronal somas (Figure 4). We clearly visualized foot like terminals of TNT-LEs on the cancer cell membrane. Elsewhere, when TNT-LEs run on the cancer cell membrane for some distance, they formed round enlargements as well as smaller granulations. Though we do not have information about their ER content, we suggest that the topology of these formations makes them strong candidates to be ERNIDs

**Figure 4.**
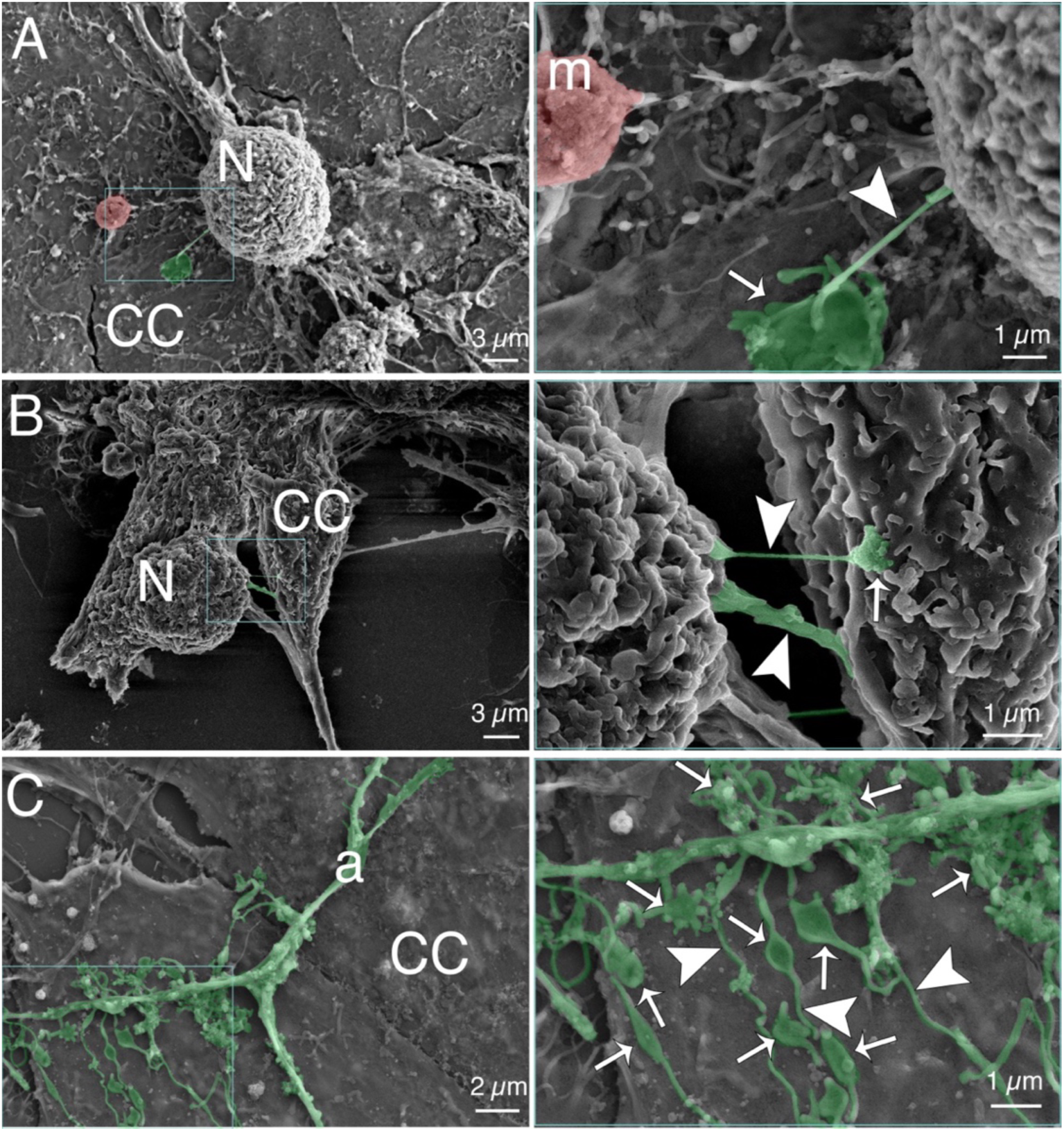
Scanning electron microscopy reveals unique morphological features of axon - cancer cell interfaces. Higher magnification images of boxed areas in the left are presented on the right and important structures are pseudo colored. **(A)** A neuron (N) sending out a TNT-LE (arrow head, green - colored) to the cancer cell (CC) underneath, with a foot – like termination (arrow); a migrasome (m, pink - colored) connected (broken) to neuron (see also Fig. S3). **(B)** Two TNT-LEs (arrow heads, green - colored) with different thickness between neuron (N) and cancer cell (CC). Arrow points to a foot like termination. (C) TNT-LEs (arrow heads, green - colored) arising from an axon (a) spread over cancer cell (CC) forming round and granular membranous structures (arrows). Note that for the purpose of clarity only some of the similar structures are pointed. Number of biological replicates (animals) / culture dish per animal / imaged area per culture dish: 3/3/5.

### Neuron-cancer cell conduits form dynamically and remodel the cancer cell surface

Finally, we performed time-lapse holotomographic microscopy (HTM), which provides high resolution three - dimensional microscopic imaging of unlabelled cells. In a representative sequence, a cancer cell lamellipodium approached an axon and formed multiple transient contacts; after lamellipodial retraction, several thin TNT-LE bridges persisted between the axon and cancer cell (Fig. 5 and Video 3). Round, highly refractile structures subsequently appeared near the contact region and accumulated across the adjacent cancer cell surface. A noteworthy observation was that during the course of establishment of TNT-LE bridges and with the emergence of round structures, the raffling activity of lamellipodia in the involved side subdued, providing a stable platform for these new formations (Fig. S2, Video 3). Correlative fluorescence-holotomographic imaging showed that bright axonal structures included ER- and mitochondria-labeled compartments (Fig. S2).

**Figure 5.**
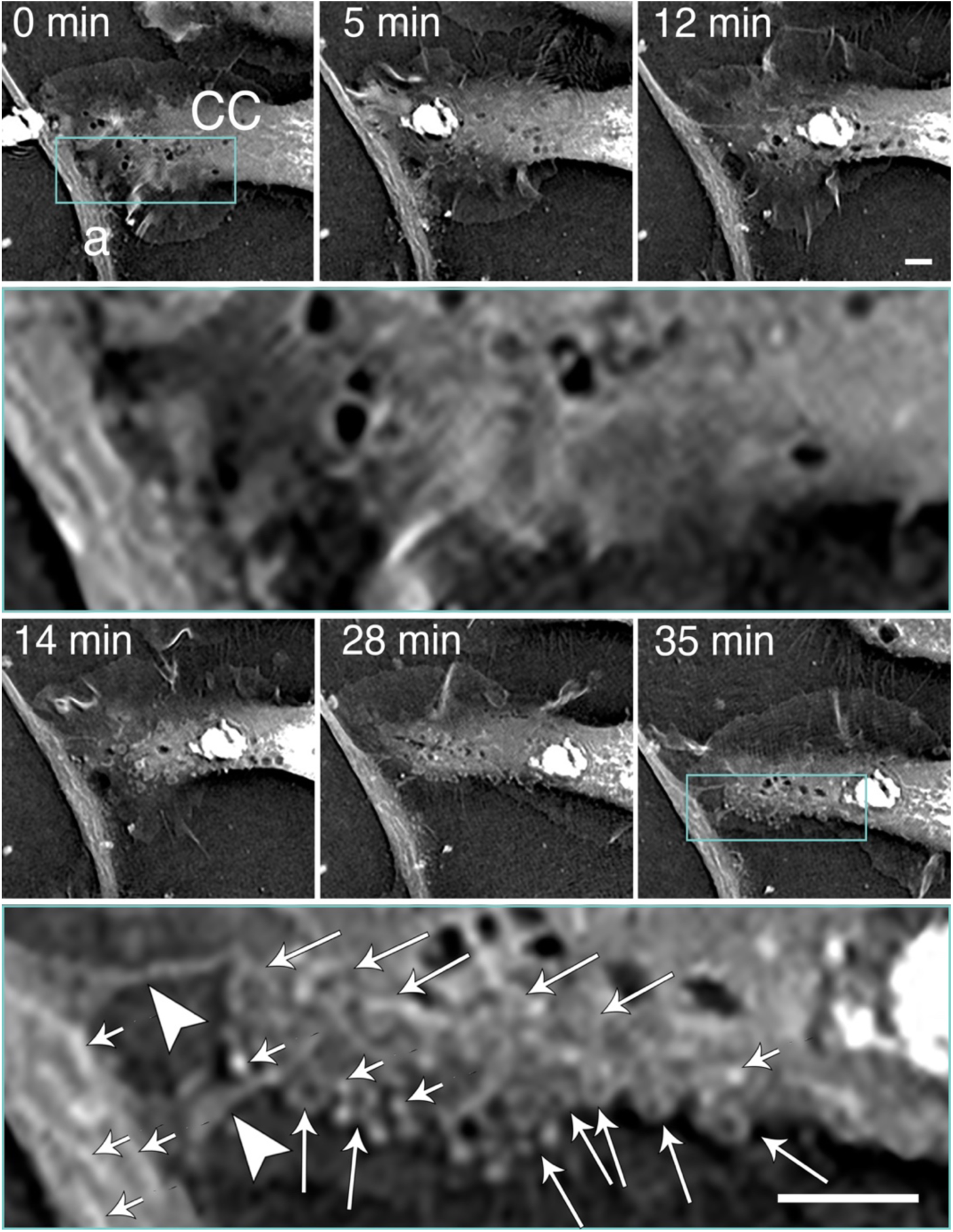
Neuron-cancer cell conduits form dynamically and remodel the cancer cell surface as revealed by time-lapse holotomographic microscopy (HTM). The second and the fourth rows are higher magnifications of boxed areas at the start and 35^th^ minutes of imaging. Arrow heads point to two TNT-LEs arising from an axon (a). Round structures emerged and progressively populated cancer cell (CC) membrane around TNT-LE connection site (long arrows). Bright refractile streaks and spots (short arrows) are likely ERs and/or mitochondria as evidenced with correlative microscopy (see Fig. S3). Note that for the purpose of clarity only some of the similar structures are pointed. Scale bars 3 µm. Number of biological replicates (animals) / culture dish per animal / imaged area per culture dish: 5/1/1. See also Fig. S3 and Video 3.

Additional electron microscopy revealed occasional mitochondria and migrasome-like structures at neuron-cancer cell interfaces (Fig. S3).

## Discussion

With very long axons, peripheral neurons have the largest cell volume in entire body. For their demanding metabolic activity, they have rich collections molecules and organelles. Since they are quiescent cells, they freely express certain oncogenes, which are in dividing cells under strict control. All these make peripheral axons very attractive for cancer cell, which is an expert of exploiting all available resources to divide and spread. Peripheral axons respond to various growth factors to grow and innervate new targets, which make them vulnerable to be lured by cancer cells. Multitude of mechanisms have been revealed about how cancer cells manipulate and use axons and new therapeutic approaches to cancer have been suggested accordingly (Jiang et al., 2026; Shin et al., 2026; Sun et al., 2026; Wang et al., 2026; Winkler et al., 2026). One of the most striking mechanisms is mitochondrion steal by cancer cell as shown both in central and peripheral tumors where TNTs act as conduits for the transfer (Hoover et al., 2025; Storevik et al., 2026). Consistent with this we have observed several mitochondria within TNT-LEs; but more importantly, we have extended this organelle affiliation to ER.

In cancer – sensory neuron co-cultures, we visualized tubular conduits of varying diameters. Classification of TNTs and similar structures are often based on their thickness and connection types (close vs. open-ended)(Cordero Cervantes and Zurzolo, 2021; Yamada et al., 2026). In this concise report, we did not attempt to identify subtypes of these conduits, instead preferred to use a general term TNT-like extensions to cover all known subspecies: filopodia, cytonemes, thin and thick TNTs and tumor microtubes (Budinger and Heneka, 2026; Cordero Cervantes and Zurzolo, 2021; Yamada et al., 2026). We detected cancer cell – ER association not only with TNT-LEs but also with axons. ERNIDs formed at such sites were larger as in the case of cap-like formations. It is noteworthy that beside TNT-LEs fused to cancer cells, we spotted some others terminating at vacuole-like invaginations as described by others (Sartori-Rupp et al., 2019). Yet, the nature of this interaction remains to be understood.

We originally defined ERNIDs depending on fluorescence confocal images of ER-tracker and GFP-tagged ER proteins CNPY2 and ATL3. High magnification SEM images suggested that ERNIDs may have multiple round membranous structures formed on the cancer cell membrane. In some STEM sections ERNIDS had profiles of round granular structures that may represent two-dimensional appearance of the same structures. Strikingly, we observed with HTM that after TNT-LE connection was established with axon, increasing number of round structures formed on the surface of cancer cell starting around TNT-LEs’ contact site, with simultaneous suppression of membrane raffling. Another interesting finding is the increased organelle transport to the axon as TNT-LEs and round structures form. We confirmed that bright regions in axons contain both ER and mitochondria in HTM images. Some of the round structures also had similar refractile patterns but whether they also represent organelles is not clear.

The existence of ER in cancer cell – neuronal conduits may be interpreted as an epiphenomenon, however high ER density at ERNIDs suggests a special formation. Advancement of ER through newly formed TNT-LEs is an energy - dependent process and ER accumulation should require further processing. These suggest existence of a specific mechanism induced to associate axonal ER to cancer cell.

Chemical synapses or pseudo synapses have been shown to form between cancer cells and peripheral axons (Fu et al., 2026; Jiang et al., 2026; Ren et al., 2025). ERNIDs can contain such structures, however, their high ER content suggests that instead of being mere presynaptic terminals, they should have more complicated functions; making them more akin to a putative extracellular organelle.

Cancer cells may benefit from neuronal ER potentially in many ways. Tubular (smooth) ER forms a continuous channel within axon, even called “neuron within neuron” (Berridge, 1998; Berridge, 2002), and it may discharge Ca^++^ to the cancer cell at ERNIDs, which will enhance their capacity for proliferation and invasion (Monteith et al., 2017). Glucose is concentrated in ER in G6P form which may potentially be served to cancer cells via ER - containing TNT-LEs (Lizak et al., 2019; Scussolini et al., 2019). Cancer cells may use lipids synthesized in tubular axonal ER to make membranes to repair and replace organelles and plasma membrane or to expand and generate new cells. While classic tubular ER lacks translation machinery and focuses strictly on lipid synthesis and calcium handling, the axonal tubular ER directly binds ribosomes and actively supports local protein synthesis (Koppers et al., 2024). This may turn ERNIDs into discrete platforms for local protein translation and processing, catering for the cancer cell.

In this preliminary study, we did not delve into the molecular or mechanistic explanation of ERNID formation. We regularly visualized mitochondria in TNT-LEs along with ER and even extracellularly. The growing interest in ER-mitochondria and other contact sites may expand into the field of cancer neuroscience (Roy et al., 2026). Presence of common mediators like CD38, which induces both extracellular and TNT-bound mitochondrial transfer and takes part in ER dynamics and potentially in TNT formation suggests the requirement of a holistic approach to cancer neuroscience research (Hayakawa et al., 2016; Marlein et al., 2019; Qin et al., 2021; Zeidler et al., 2022).

A final remark is about the migrasome-like structures we visualized with STEM an SEM. Migrasomes are relatively large extracellular organelles, typically ∼0.5–3 μm, generated on retraction fibers of migrating cells. They can carry proteins, RNAs, lipids, signaling molecules, and even damaged mitochondria, and can release their contents through migracytosis.This is particularly intriguing because a recent synthesis of the field explicitly places TNTs and migrasomes as complementary communication mechanisms: TNTs provide direct cytoplasmic exchange, whereas migrasomes contribute to spatial distribution of signaling molecules in the tumor microenvironment (Li et al., 2026; Liu et al., 2026). We have imaged a migrasome-like body captured between a neuron and cancer cell with STEM and another one next to a neuron with SEM; yet we have no evidence to tell the direction of the transfer. However, in either case the cancer cells are likely to be benefited by receiving neuronal material or delivering instruction packages to neurons to guide them to enhance tumor growth.

## Limitations and Conclusions

The study has several limitations. The experiments were performed in vitro with one murine breast cancer cell line, the molecular machinery that generates the conduits was not tested, and the functional consequences for cancer cells remain unknown. We also did not determine whether ER-containing contacts occur in tumors in vivo. Nevertheless, the convergence of complementary imaging approaches establishes a previously unrecognized ER-rich axon-cancer cell interface and provides a tractable system for mechanistic studies of direct neuron-tumor communication.

## Materials and Methods

### Lentiviral vector construction and production

Coding sequences of mouse ATL3 and CNPY2 were amplified from mouse cDNA and cloned upstream of enhanced GFP in the third-generation pLV-eGFP lentiviral transfer vector (Addgene plasmid 36083), generating ATL3-eGFP and CNPY2-eGFP constructs. Assembly was performed using NEBuilder HiFi DNA Assembly (New England Biolabs), and insert orientation and sequence integrity were confirmed by Sanger sequencing. The pLV-CMV-mCherry transfer vector was used to generate mCherry-expressing EMT6 cells. Lentiviral particles were produced in HEK293T cells by cotransfection of the transfer vector with third-generation helper plasmids encoding Gag/Pol (pMDL), Rev (pRev), and VSV-G, using polyethylenimine (Polysciences, 23966-100) at a DNA:PEI ratio of 1:4. Supernatants were collected 72 h after transfection. pLV-CMV-mCherry supernatant was passed through a 0.45-µm filter and used directly. ATL3-eGFP and CNPY2-eGFP particles were concentrated through a 20% (wt/vol) sucrose cushion by centrifugation at 18,000 × g for 4 h at 4°C, resuspended in 100 µl PBS, aliquoted, and stored at -80°C.

### Generation of stable mCherry-expressing EMT6 cells

EMT6 murine breast cancer cells were transduced with pLV-CMV-mCherry lentiviral supernatant. mCherry expression was confirmed by fluorescence microscopy, and mCherry-positive cells were isolated by fluorescence-activated cell sorting using a BD Influx cell sorter (BD Biosciences). Sorted cells were expanded and used for co-culture experiments.

### Primary DRG neuron culture and co-culture

The use of animals in this study was approved by Istanbul Medipol University Animal Experimentation Ethical Committee (approval no. 29, 22 May 2024). Maximum care was paid to reduce the minimum number of animals and to minimize suffering. All procedures were conducted according to institutional guidelines which are in compliance with European Council Directive 2010/63/EU. Breeding colonies of the mice were obtained from The Jackson Laboratory (US) and bred at Istanbul Medipol University Experimental Animal Facility. Primary dorsal root ganglia (DRGs) were isolated from adult TauEGFP knock-in mice (The Jackson Laboratory, stock no: 004779, RRID:IMSR_JAX:004779) (Tucker et al., 2001) or wild type C57 strains.After euthanasia by gradual CO_2_ inhalation. DRGs were dissected into cold RPMI-1640, stripped of attached nerve fibers, and transferred to Neurobasal-A medium supplemented with 2% B27, 2 mM GlutaMAX-I, 100 U/ml penicillin, 100 µg/ml streptomycin, 250 ng/ml amphotericin B, and 100 U/ml collagenase. Tissue was incubated for 50 min at 37°C and 5% CO_2_, washed three times with HBSS, and digested with 1 mg/ml trypsin for 15 min. DRGs were mechanically dissociated, treated with 50 µg/ml DNase I for 30 min at 37°C, centrifuged, and resuspended in Neurobasal-A containing 10% FBS and 30 µg/ml trypsin inhibitor. Neurons were enriched using a three-layer Percoll gradient (60%, 35%, and 10%) and collected from the 35% layer after centrifugation at 1,700 rpm for 20 min at 4°C. Purified neurons were plated on 35-mm glass-bottom dishes coated with poly-L-lysine (1.8 µg/cm2) and laminin (40 ng/mm2). After 2 h, unattached cells and debris were removed and fresh Neurobasal-A medium with the supplements was added. mCherry-positive EMT6 cells were added directly to DRG cultures, and co-cultures were maintained at 37°C and 5% CO_2_.

### ER labeling and confocal microscopy

For ER-Tracker experiments, DRG cultures were incubated with 0.5 µl ER-Tracker Blue-White DPX (Thermo Fisher Scientific, E12353) for 24 h and medium was changed with fresh Neurobasal-A (with the supplements) after washing three times with sterile PBS. Following another 24 hrs incubation, EMT6-mCherry breast cancer cells were added to the cultures.

For genetically encoded ER labeling, DRG neurons were transduced with ATL3-eGFP or CNPY2-eGFP viral vector on the first day of culture and incubated for 48 h. Then medium was changed with fresh Neurobasal-A (with the supplements) after washing three times with sterile PBS before adding EMT6-mCherry cells.

On DIV3, co-cultures were imaged using a Zeiss LSM 800 laser-scanning confocal microscope with a 40× oil-immersion objective (N.A. 1.30) employing Airyscan utilty. For blue, green and red fluorescence imaging 405, 488 and 561 laser lines were used, respectively. Image acquisiton and processing were performed with Zen Black and Zen Blue software suits.

### Scanning electron microscopy (SEM)

On DIV3, some co-cultures were fixed in Karnovsky’s fixative for 1 h at room temperature, washed in 0.15 M sodium cacodylate buffer, and postfixed in 2% osmium tetroxide for 30 min. Samples were dehydrated through graded ethanol, exchanged into acetone, critical-point dried with liquid CO_2_ (Leica EM CPD300), mounted on aluminum stubs, sputter-coated with gold-palladium for 10 s (Leica EM ACE200), and imaged using a GeminiSEM 500 field-emission scanning electron microscope at 5 kV with an SE2 detector and a working distance of 6.7 mm.

### Serial-section scanning transmission electron microscopy (STEM)

On DIV3, some co-cultures were fixed in Karnovsky’s fixative, washed in 0.15 M sodium cacodylate buffer, and processed with reduced osmium (2% osmium tetroxide and 1.5% potassium ferrocyanide), 0.5% thiocarbohydrazide, a second 2% osmium step, 2% aqueous uranyl acetate overnight at 4°C, and lead aspartate for 20 min at 60°C. After graded ethanol dehydration, samples were infiltrated with Epon resin and polymerized overnight at 60°C. Consecutive 100-nm sections were cut with a diamond knife on a Leica EM UC7 ultramicrotome, collected on glass coverslips, sputter-coated with gold-palladium, and imaged on a GeminiSEM 500 at 5 kV using a backscattered-electron detector at a working distance of 6.2 mm.

### Time-lapse holotomographic microscopy

On DIV2 of DRG cultures, EMT6 cells were added to the dish and without waiting, preparations were placed on a holotomographic microscope (Nanolive 3D Cell Explorer) equipped with a stage-top incubator maintained at 37°C and 5% CO_2_. Images were acquired every minute for 12 hours with a 60× objective (N.A. 0.8) using STEVE software. Three-dimensional 96 image stacks were processed in ImageJ. For correlative imaging, ER-Tracker and MitoTracker fluorescence was aligned with holotomographic images.

### Statistics and reproducibility

This study is descriptive and no statistical hypothesis tests were performed. Numbers of biological replicates, cultures and imaged fields are provided in related figure legends.

## SUPPLEMENTAL MATERIAL

**Figure S1:**
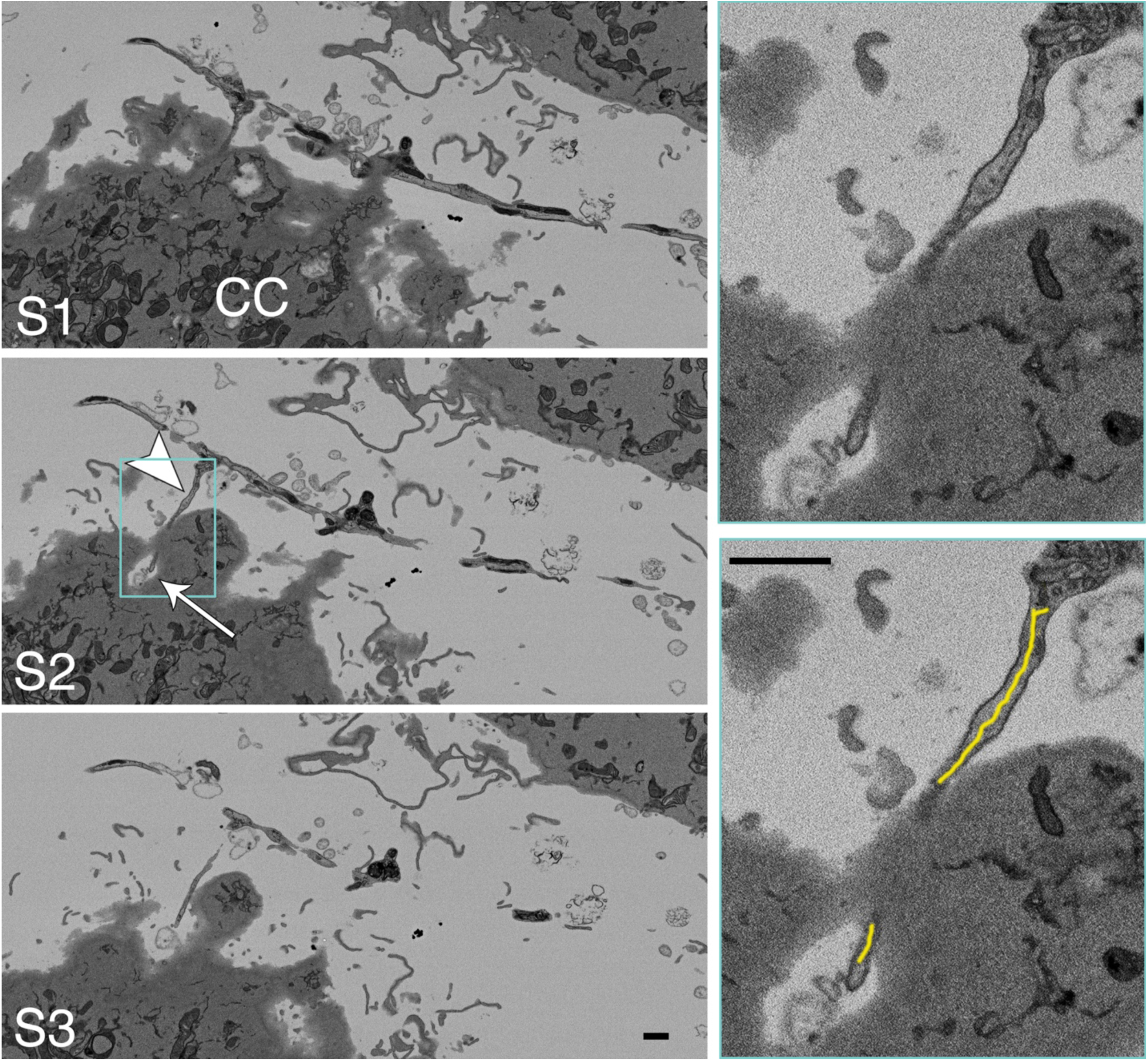
Some TNT-like extensions (TNT-LEs) terminate in vacuole-like structures in cancer cells. Samples (S1-3) from serial sections of a cancer cell (CC) receiving an incoming TNT-LE (arrow head) into a round cytoplasmic cavity (arrow). Magnified view of the boxed area is presented on the right. The TNT-LE contains ER (yellow trace). Scale bars 1 µm. Number of biological replicates (animals) / culture dish per animal / sectioned and imaged area per culture dish: 3/3/1.

**Figure S2:**
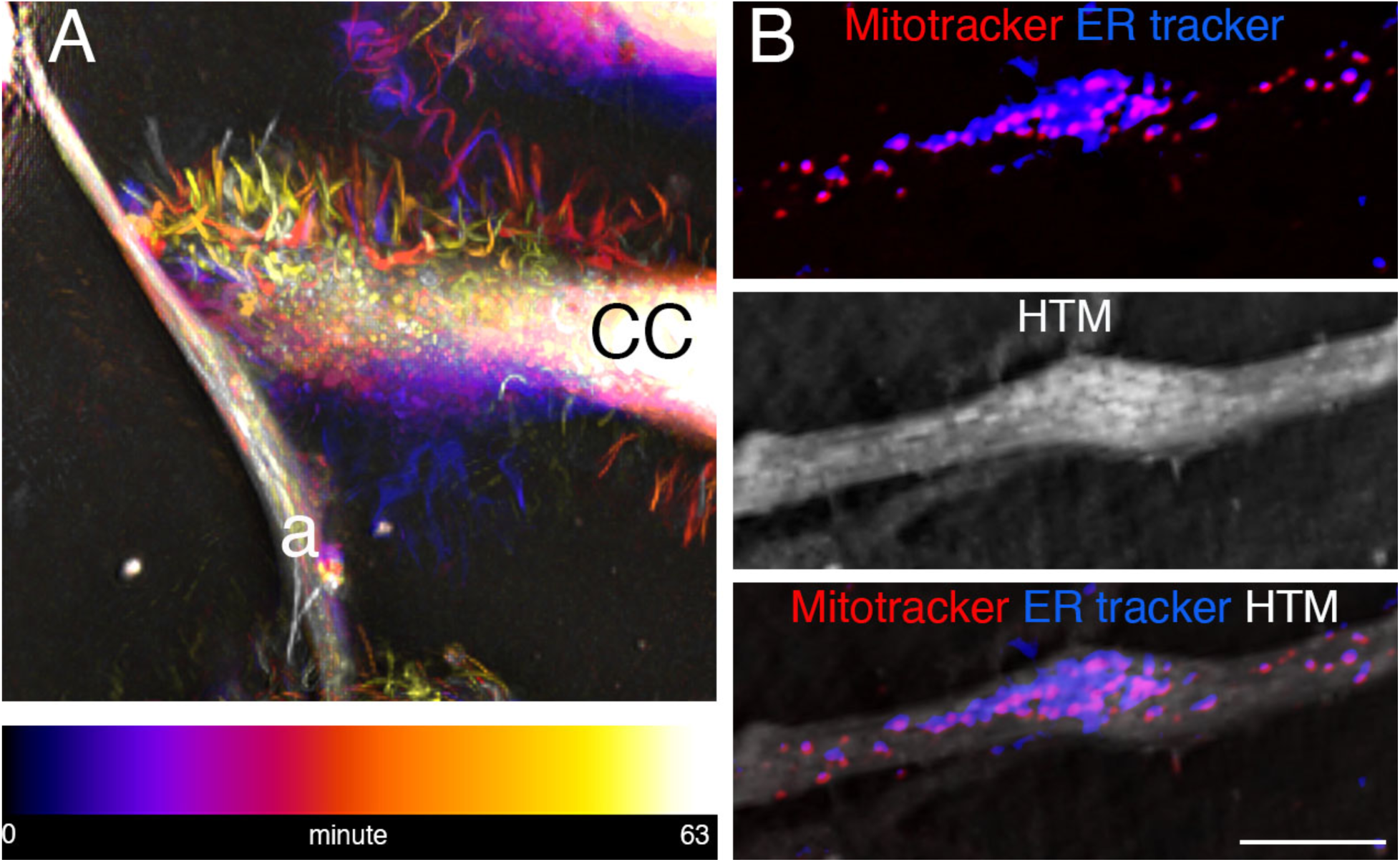
Holotomographic microscopy (HTM) reveals temporal and spatial features of axon – cancer cell interactions. **(A)** Time-coded activities during axon (a)-cancer cell (CC) interactions during 63 min incubation period **(B)** Correlative confocal –HTM imaging reveals identity of bright structures as ERs and mitochondria. Scale bar 5 µm. Number of biological replicates (animals) / culture dish per animal / imaged area per culture dish: 5/1/1. See also Fig. 5 and Video 3.

**Figure S3:**
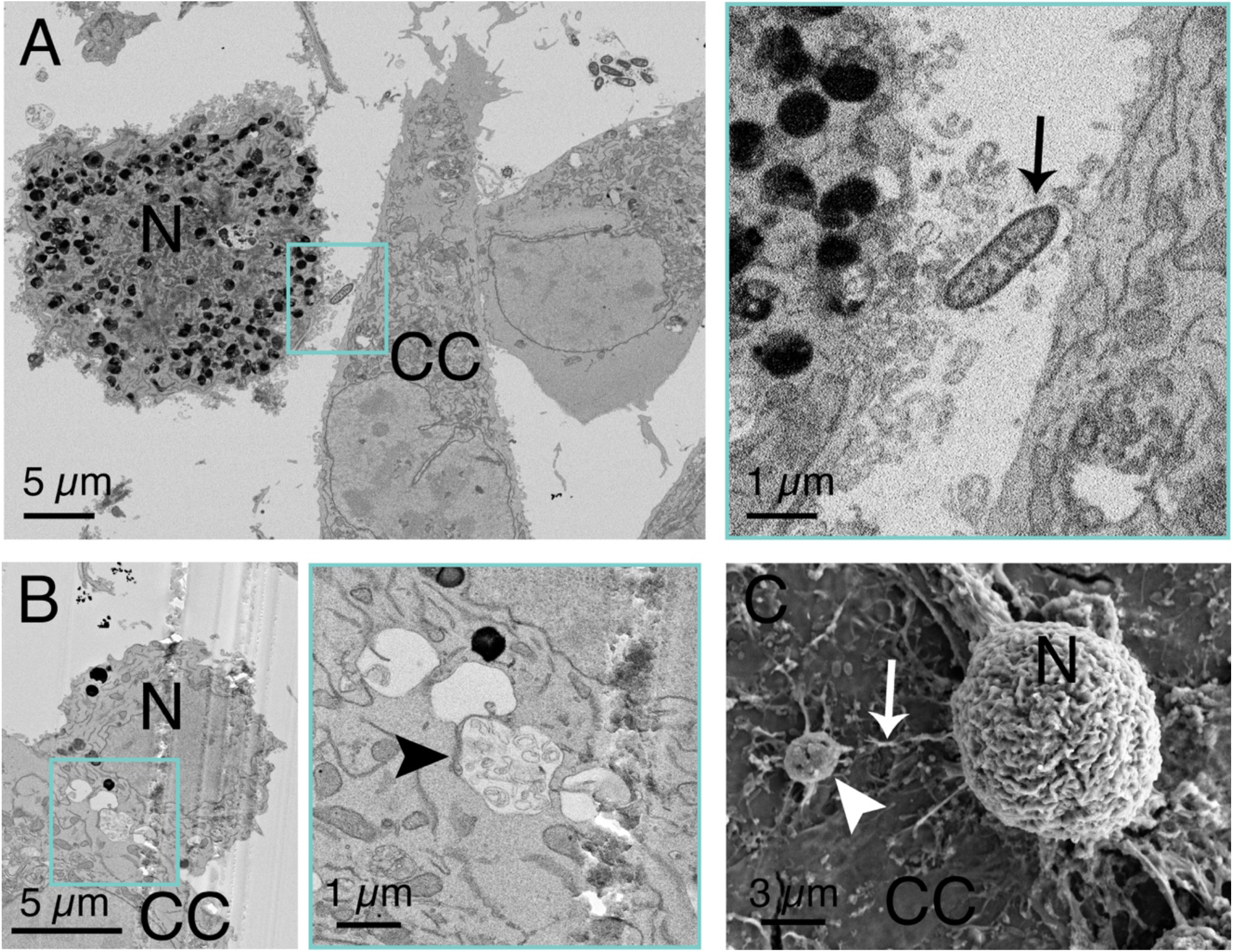
Electron microscopy reveals a possible mitochondrion transport and migrasomes in neuron – cancer cell co-cultures. **(A)** A mitochondrion (arrow) between neuron (N) and cancer cell (CC), which may be in transport. **(B)** A migrasome (arrow head) between cancer cell (CC) and neuron (N). **(C)** A migrasome (arrow head) with a TNT-like connection (arrow) to neuron (N) lying on cancer cell (CC); the connection was probably broken during tissue processing. Number of biological replicates (animals) / culture dish per animal / sectioned and imaged area per culture dish: 3/3/1.

### Video Captions

**Video 1.** Connection of a CNPY2 expressing neuron (green) with TNT-LE to cancer cell (red) with an ER rich termination. See also Figure 2.

**Video 2.** Connection of an ATL3 expressing neuron (green) with TNT-LE to cancer cell (red) with ER rich terminal domains. See also Figure 2.

**Video 3.** Time lapse recording of interaction of an axon and a cancer cell resulting in formation of TNT-LEs and round membranous structures on the cancer cell membrane.

